# Control of division and microtubule dynamics in *Chlamydomonas* by cyclin B/CDKB1 and the anaphase-promoting complex

**DOI:** 10.1101/2021.06.29.450376

**Authors:** Kresti Pecani, Kristi Lieberman, Natsumi Tajima-Shirasaki, Masayuki Onishi, Frederick R. Cross

## Abstract

In yeast and animals, cyclin B binds and activates the cyclin-dependent kinase (‘CDK’) CDK1 to drive entry into mitosis. We show that CYCB1, the sole cyclin B in *Chlamydomonas*, activates the plant-specific CDKB1 rather than the CDK1 ortholog CDKA1. Time-lapse microscopy shows that CYCB1 is synthesized before each division in the multiple fission cycle, then is rapidly degraded 3-5 minutes before division occurs. CYCB1 degradation is dependent on the anaphase-promoting complex (APC). Like CYCB1, CDKB1 is not synthesized until late G1; however, CDKB1 is not degraded with each division within the multiple fission cycle. The microtubule plus-end-binding protein EB1 labeled with mNeonGreen (EB1-NG) allowed detection of mitotic events in live cells. The earliest detectable step in mitosis, splitting of polar EB1-NG signal into two foci, likely associated with future spindle poles, was dependent on CYCB1. CYCB1-GFP localized close to these foci immediately before spindle formation. Spindle breakdown, cleavage furrow formation and accumulation of EB1 in the furrow were dependent on the APC. In interphase, rapidly growing microtubules are marked by ‘comets’ of EB1; comets are absent in the absence of APC function. Thus CYCB1/CDKB1 and the APC mitosis modulate microtubule dynamics while regulating mitotic progression.

## INTRODUCTION

Control of the eukaryotic cell cycle has been extensively characterized in animals and yeast (Opisthokonts), but less is known in other eukaryotes, including the plant kingdom, which diverged from Opisthokonts early in evolution (Rogozin et al., 2009). *Chlamydomonas reinhardtii* is a microbial member of the plant kingdom with unique advantages for studying basic cell biology compared to land plants: mostly single-copy genes, a simple unicellular life cycle, and facile Mendelian and molecular genetics. Genetic experiments have shown that as in yeast and animals, cyclin B is essential for cell division, and the anaphase promoting complex (APC) is essential for anaphase and exit from mitosis (Tulin & Cross, 2014, Atkins & Cross, 2018). We showed previously that the plant-specific CDKB1 is the essential CDK for mitotic entry, rather than CDK1/CDKA1 as in yeast and animals, and CDKB1-associated kinase activity was genetically dependent on CYCB1 (Atkins & Cross, 2018). Here, we used tagged transgenes to confirm specific CYCB1-CDKB1 interaction. We developed methods for long-term time-lapse fluorescent microscopy of single cells, and measured accumulation and degradation of CYCB1 and CDKB1 through cycles of multiple fission. In addition, we used mNeonGreen-tagged EB1 (microtubule plus-end-binding protein) in WT and mutants to understand genetic requirements for microtubule dynamics and individual steps in mitotic progression.

## RESULTS

### CYCB1 interacts with CDKB1

B-type cyclins are key regulators of the cell cycle in animals and in yeast (Morgan, 2007). *Chlamydomonas* has a single essential cyclin B gene, *CYCB1* (Atkins & Cross, 2018). We constructed a *CYCB1-GFP* fusion under control of the *CYCB1* promoter, and identified transgene transformants that rescued ts-lethality of *cycb1-5*. We chose a transformant with a single GFP-containing locus that efficiently rescued *cycb1-5* in tetrad analysis. Immunoblotting with anti-GFP revealed a single CYCB1-GFP band, expressed specifically in dividing cells in partially synchronized cultures (Fig. 1). Inactivation of either the APC or of CDKB1 (using *cdc27-6* or *cdkb1-1* mutations, respectively) prevented degradation of CYCB1-GFP at late timepoints (Fig. 1).

**Figure 1.**
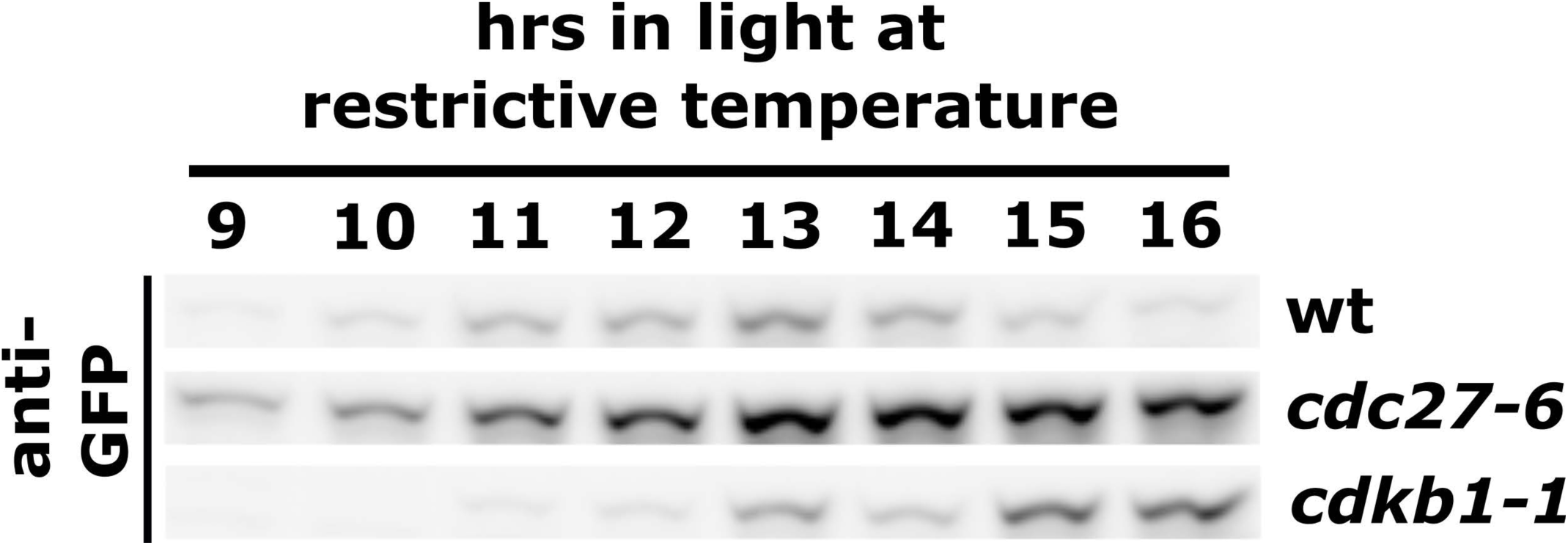
Detection of CYCB1-GFP in wt, cdc27-6, or cdkb1-1 backgrounds by immunoblotting. Anti-GFP immunoblotting of cells with temperature-sensitive mutations cdc27-6 or cdkb1-1. Cells were placed at restrictive temperature and collected after the indicated number of hours. All strains had temperature-sensitive cycb1-5 rescued by CYCB1-GFP transgene.

*In vitro* protein kinase activity toward histone H1 co-immunoprecipitated with CYCB1-GFP (Fig. 2A). This activity was eliminated in *cdkb1-1 CYCB1-GFP* cells despite the presence of CYCB1-GFP protein (Fig. 2A). Reciprocally, the CDKB1-associated kinase activity was genetically dependent on *CYCB1* (Atkins & Cross, 2018). Kinase activity was increased in *cdc27-6 CYCB1-GFP* cells (Fig. 2A); similarly. CDKB1-associated kinase activity was increased in *cdc27-6 CDKB1-mCherry* (Atkins & Cross, 2018).

**Figure 2.**
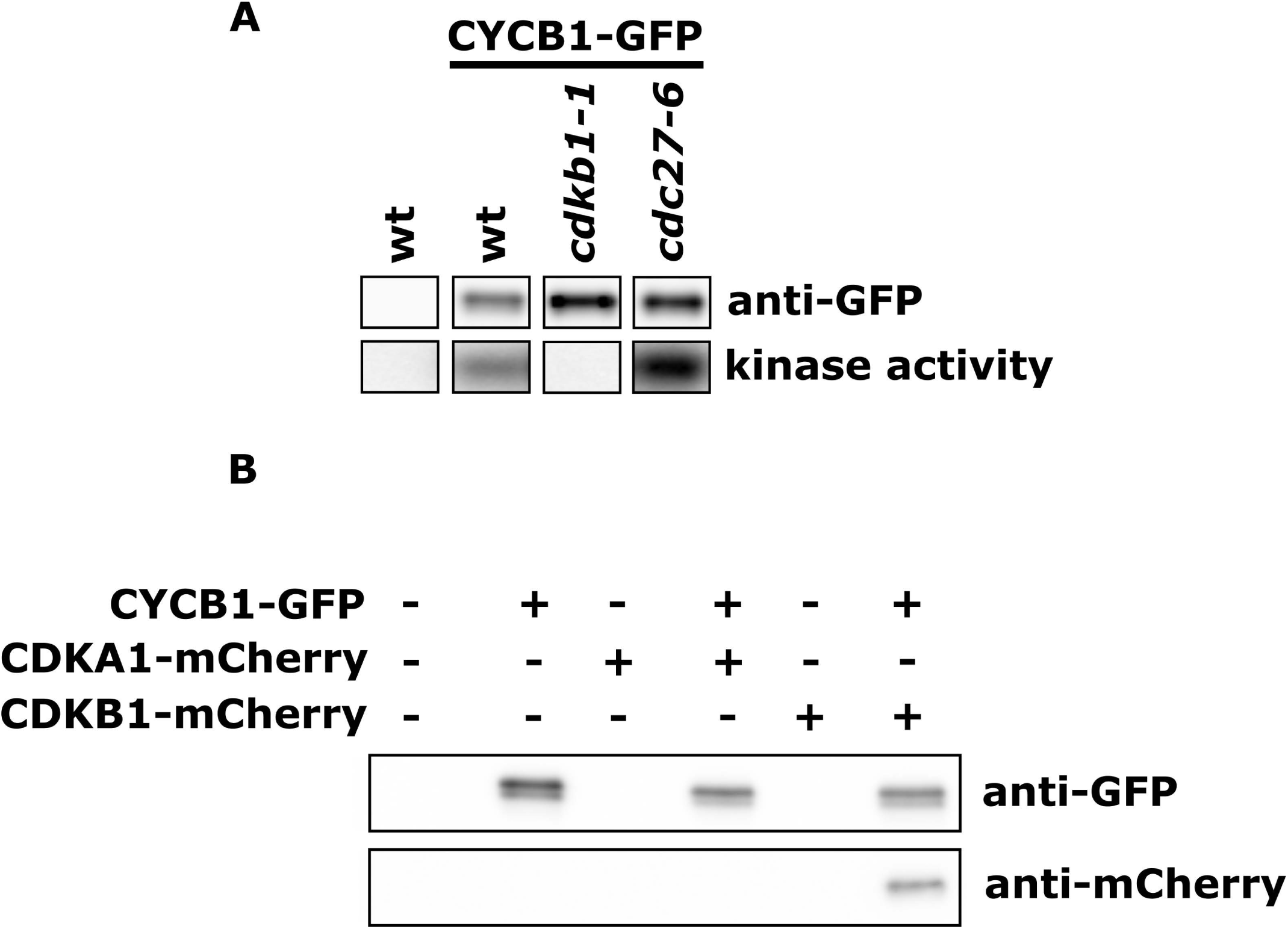
Detection of CYCB1-GFP binding partners and kinase activity by co-immunoprecipitation. A: Anti-GFP immunoblotting of CYCB1-GFP pull-down in untagged control (‘wt’), wt, cdkb1-1, or cdc27-6 backgrounds (top row). Kinase activity co-immunoprecipitated with CYCB1-GFP in untagged, wt, cdkb1-1, or cdc27-6 backgrounds (bottom row). All strains except for untagged wt control on left had temperature-sensitive cycb1-5 rescued by CYCB1-GFP transgene. B: Detection of CDKA1-mCherry or CDKB1-mCherry as possible binding partners of CYCB1-GFP. Strains with CYCB1-GFP and CDKA1-mCherry or CDKB1-mCherry (and wt, CDKA1- mCherry or CDKB1-mCherry alone) were immunoprecipitated with anti-GFP. Immunoblotting was then done with anti-GFP or anti-mCherry.

Specificity of interactions between cyclins and CDKs in *Arabidopsis* has been inconclusive. Comprehensive proteomics with tagged proteins showed that cyclin B bound specifically to CDKB and not CDKA (Van Leene et al., 2010); however, Boruc et al., 2010 showed by binary interaction assays that CDKB and CDKA both have the capacity to bind CYCBs and CYCAs. We constructed *Chlamydomonas* CYCB1-GFP strains co-expressing CDKA1-mCherry or CDKB1-mCherry. Anti-GFP immunoprecipitation specifically co-precipitated CDKB1-mCherry but not CDKA1-mCherry (Fig. 2B). This finding contrasts with the specificity of Opisthokont cyclin B to the CDKA1 ortholog CDK1.

Overall, these data suggest that early in the evolution of the plant lineage, the plant-specific CDKB1 took over the role of inducing mitotic progression in response to cyclin B accumulation. Experiments in the microalga Ostreococcus also support this idea (Corellou et al., 2005).

### Cyclin B accumulation and degradation through multiple fission

*Chlamydomonas* exhibits a pattern of cell division called ‘multiple fission’ (Cross & Umen, 2015). Newborn cells are small, and can grow over a 10-12 hr period to >10-fold starting size without DNA synthesis or cell division. Cells then resorb flagella and undergo multiple rapid cell divisions: complete rounds of DNA replication, nuclear division and cytokinesis, all within the mother cell wall, until progeny cells have divided to approximately their starting size (Cross & Umen, 2015). Divisions require ∼30 min and are highly synchronous synchronous among the descendants of a single cell, which are retained within within the mother cell wall until hatching occurs after the terminal cell division (Cross & Umen, 2015).

The Western blot data above showed approximate restriction of cyclin B accumulation to the multiple fission period. However, synchrony is not good enough to resolve individual divisions in bulk culture, preventing determination of whether cyclin B was stable throughout the period of multiple fissions, or was degraded in each division and then resynthesized.

To solve this problem, we developed methods for long-term time-lapse fluorescence microscopy of *Chlamydomonas*. This required maintenance of tight temperature regulation of cells being imaged, preventing cells swimming out of the field of view, providing light for photosynthesis between image acquisitions, and computational subtraction of autofluorescence from chloroplasts, which otherwise swamped the CYCB1-GFP signal. Our solutions included small, individually-sealed acrylic chambers filled with TAP/agarose, each containing a different cell population; overhead illumination provided by small LEDs, which were programmed to turn on after image acquisition was complete, and turn off before the start of the next frame; and computational deconvolution to eliminate contribution of chloroplast autofluorescence. See time lapse microscopy method #1 in Methods section for complete details.

CYCB1-GFP signal was first detectable ∼0.5-1 hr before the first division (Fig. 3; Supp. Video 1); signal increased steadily for approximately 20 min. We then observed sharp reduction to near-background levels of the signal 3-6 min (1-2 frames) before cell division (scored by formation of a cleavage furrow in a concurrent brightfield image; arrows in Fig. 3, Supp. Video 1). From the shape and position of the signal we assume CYCB1-GFP is nuclear-localized.

**Figure 3.**
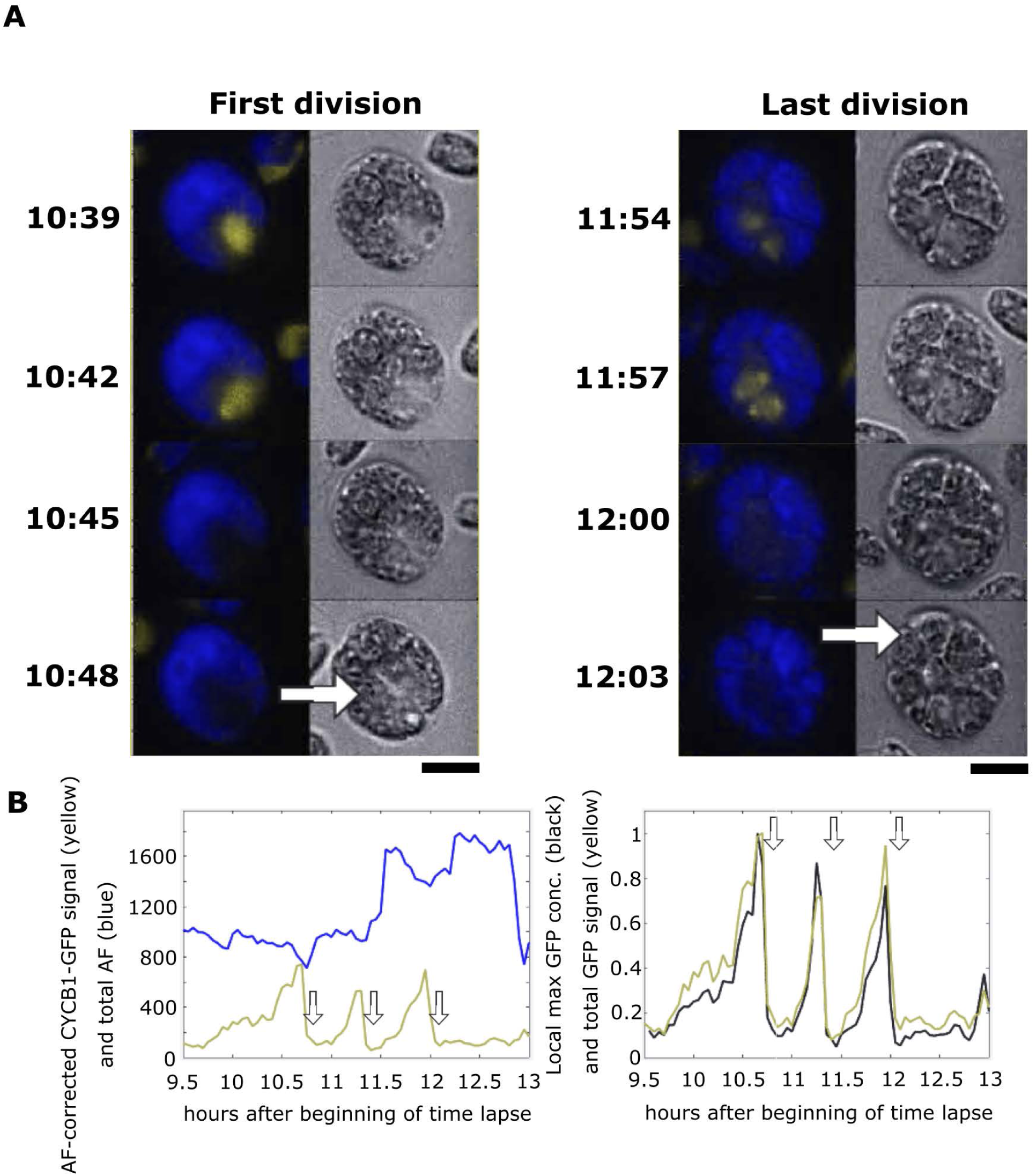
Live cell time lapse microscopy of CYCB1-GFP. (A) Time lapse images of CYCB1-GFP cells. Each cell has a brightfield image (right), and a composite of chloroplast autofluorescence in blue and CYCB1-GFP signal in yellow (left). Time indicated on top of each strip is hours and minutes after beginning of time lapse. The indicated time corresponds to the top cell in each strip. Each subsequent cell going down is from an image captured every 3 minutes (time from plating indicated). Arrows indicate new cleavage furrow formation detected in brightfield. The imaged cell went through three divisions; frames surrounding the first and last divisions are shown. Scale bar: 5 microns. (B) left: quantification of CYCB1-GFP signal deconvolved from chloroplast autofluorescence (yellow line), and chloroplast autofluorescence (blue line). Arrows: correspond to cleavage furrow formation. Right: Yellow trace: CYCB1-GFP total signal over the cell. Black: a minimal convex hull was computed that contained 50% of the CYCB1-GFP signal, and the concentration (signal/area) computed, showing that local concentration and total cellular amount of CYCB1-GFP tracked closely through divisions. MATLAB code for calculating the convex hull available on request.

From multiple movies, we estimate a half-life of nuclear CYCB1-GFP of approximately 3-5 minutes, specifically during an interval of ∼5-10 min preceding cell division. CYCB1-GFP then reaccumulates, but only in cells destined to undergo an additional division cycle. This indicates that the ‘decision’ to divide is upstream of CYCB1 accumulation. We don’t have an estimate for the half-life of CYCB1-GFP during the reaccumulation phases, but the protein accumulates in a linear fashion for at least 0.5 hr, suggesting a half-life at least this long.

Newly accumulated CYCB1-GFP in later division cycles is sometimes clearly separated into 2 (2^nd^ division) or 4 (3^rd^ division) foci, which we presume corresponds to separate accumulation in different daughter nuclei. This is not always clearly observed; we don’t know if this is due to specifics of nuclear localization within daughter cells or to complexity of the multiply divided cells’ geometry observed at a single focal plane.

### Cyclin B proteolysis is dependent on APC and on CDKB1

APC-dependent ubiquitination and proteolysis is frequently dependent on a ‘destruction box’ consensus sequence in the target protein (He et al., 2013). CYCB1 contains a consensus destruction box (Atkins & Cross, 2018). Using *cdc27-6*, a tight temperature-sensitive allele of a core APC subunit (Atkins & Cross, 2018), we found that CYCB1-GFP proteolysis was dependent on the APC (Fig. 4): CYCB1-GFP levels were low at early times, and rose at similar times to WT, but unlike in WT, no precipitous degradation was observed even after many hours.

**Figure 4.**
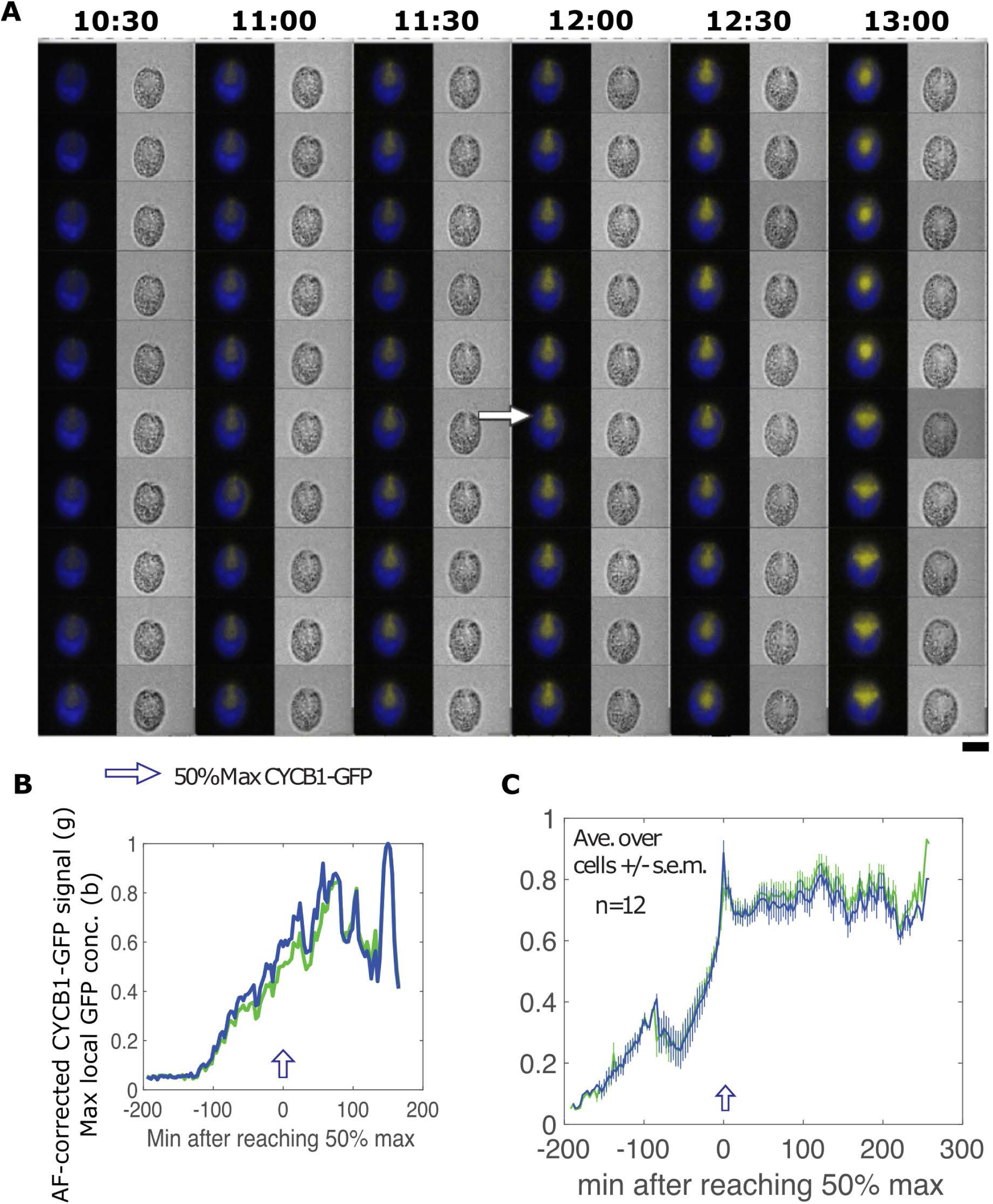
Live cell time lapse of CYCB1-GFP in a *cdc27-6* background. (A) Each cell has a brightfield image (right), and a composite of chloroplast autofluorescence in blue and CYCB1-GFP signal in yellow (left). Time indicated on top of each strip is hours and minutes after beginning of time lapse. The indicated time corresponds to the top cell in each strip. Each subsequent cell going down is from an image captured every 3 minutes. Scale bar: 5 microns. (B) Green: deconvolved total GFP signal in cell shown in A; blue: concentration estimated as in Fig. 3. (C) The same plots for the average and s.e.m. of 12 cells. All traces adjusted to a maximum signal of 1 before averaging.

We also found that CYCB1-GFP levels remained high in a *cdkb1-1* background (Supp. Fig. 1, Supp. Video 2). This observation could be explained in two ways: (1) Degradation might be restricted to CYCB1 in a complex with CDKB1. (2) CYCB1-CDKB1 might be required to activate the APC. The former may be unlikely since in other organisms, APC-dependent degradation generally transfers with the destruction box, even if appended to reporters (Glotzer et al., 1991). The latter mechanism could be consistent with results in animal cells, APC-Cdc20 activation is dependent on cyclin B-Cdk1 phosphorylation of APC subunits (Zhang et al., 2016). The mechanism is complex: recruitment of Cks1-CDK-cyclin to a disordered region of APC3, promoting phosphorylation of a segment of APC1 that occludes the Cdc20 binding site; phosphorylated APC1 does not occlude the site and Cdc20 is recruited (Zhang et al., 2016). The regions and phospho-sites in human APC3 and APC1 identified as critical for this mechanism align poorly or not at all to the *Chlamydomonas* homologs, so if a similar mechanism is operating, it is working with divergent sequences. We have observed complete synthetic lethality at permissive temperature in tetrad analysis between temperature-sensitive mutations in *CDC20* and *CKS1* (Breker et al., 2018; unpublished data), suggesting some collaboration between CDC20 and CKS1, but we have no information specifically connecting this to CDC20 activation by CDK beyond the ability of CKS1 to bind CDKB.

### Is Cyclin B degradation essential?

In yeast and animals, cyclin B degradation is essential for completion of cytokinesis and for initiation of a new round of DNA replication (Murray & Kirschner, 1989; Wäsch & Cross, 2002). In yeast, this requirement for cyclin B degradation is specific to the Clb2 B-type cyclin; mitotic exit proceeds even without the destruction of another B-type cyclin, Clb3, and the degradation of Clb3 is not essential for viability (Pecani & Cross, 2016). In *Nicotiana tabacum*, expression of non-degradable CYCB1 leads to endomitosis with failed cytokinesis (Weingartner et al., 2004). We therefore tested whether destruction of CYCB1 is essential in *Chlamydomonas*.

We constructed a *CYCB1-db-GFP* transgene with the destruction box deleted, and transformed it into a *cycb1-5* temperature-sensitive strain in parallel with wild-type *CYCB1-GFP*, selecting at 33 degrees for rescue of *cycb1-5.* In three independent experiments, each with hundreds of rescue events by WT *CYCB1-GFP*, we obtained no rescue upon electroporation with similar amounts of *CYCB1-db-GFP* (data not shown). This result is consistent with lethality of *CYCB1-db*. We cannot rule out the possibility that the CYCB1 destruction box is required for positive function of CYCB1; this has not been observed in other systems, however, and the destruction box is far from the cyclin regions responsible for CDK activation and substrate targeting. At any rate, these results indicate that the destruction box in CYCB1 is essential for its function.

### Regulation of CDKB1

We reported previously that CDKB1-mCherry accumulated in the nucleus of cells during the multiple fission period (Atkins & Cross, 2018); however, we were unable to resolve whether CDKB1-mCherry was degraded and then resynthesized in each cell cycle. We constructed a CDKB1- Venus transgene, and used it to rescue a *cdkb1-1* strain. In time lapse microscopy of rescued cells, we observed CDKB1-Venus accumulation tightly specific to the period of the multiple fission cycle, consistent with previous results (Atkins and Cross, 2018). However, there was no loss of total CDKB1-Venus signal within the individual divisions, unlike the behavior of CYCB1-GFP (Fig. 5).

**Figure 5.**
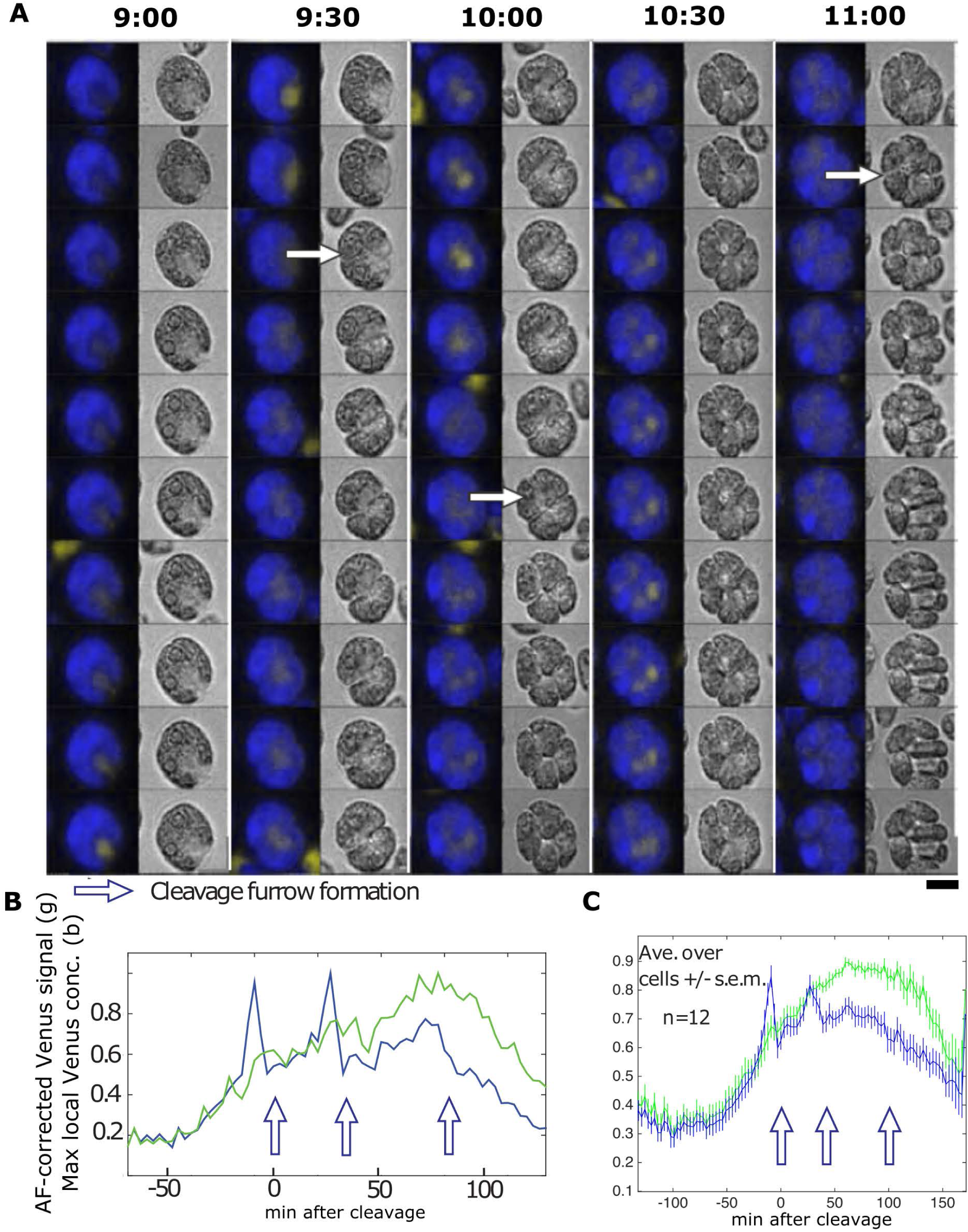
Live cell time lapse of CDKB1-Venus cells. (A) Brightfield image (right), and a composite of chloroplast autofluorescence in blue and CDKB1-Venus signal in yellow (left). Time indicated on top of each strip is hours and minutes after beginning of time lapse. The indicated time corresponds to the top cell in each strip. Each subsequent cell going down is from an image captured every 3 minutes. Arrows: cleavage furrow formation. Scale bar: 5 microns. (B) Green line: total YFP signal; blue line: estimated concentration (YFP signal per area) in the minimal convex hull calculated to contain 50% of total signal. (C) The same plots for the average and s.e.m. of 12 cells. All traces adjusted to a maximum signal of 1 before averaging. In B and C, note reproducible peak of concentration of CDKB1-GFP 1-2 frames before cleavage furrow formation.

Although CDKB1-Venus signal quantified over the entire cell remained high through multiple division cycles, the local intensity of the nuclear signal varied through the cell cycle, reaching a peak about 6 min before division (Fig. 5). The timing of more intense CDBK1-Venus localization approximately corresponds to the timing of CYCB1 accumulation. We speculate that efficient nuclear localization of CDKB1 may require CYCB1.

After completion of the terminal cell division, CDKB1-Venus remained diffuse and disappeared over the succeeding ∼1 hr, suggesting that CDKB1 degradation might be dependent on exit from the multiple fission period.

### Live-cell imaging with EB1-mNeonGreen reveals regulation of microtubule and spindle dynamics

Cytoplasmic microtubules in *Chlamydomonas* are of two types. There are very stable ‘rootlets’ forming a cruciate structure centered on the basal bodies, containing acetylated tubulin (Ehler et al., 1995; Janke & Montagnac, 2017). In addition, there are unacetylated and highly dynamic ‘cytoplasmic microtubules extending from the vicinity of the basal bodies and rootlets and forming a cup-shaped pattern with the basal bodies as the base’ (Ehler et al., 1995). The plus-end-binding protein EB1-mNeonGreen (EB1-NG) is located in one or two anterior spots at or near to the flagellar basal bodies (Harris et al., 2016; Onishi et al., 2020), and moving EB1-NG ‘comets’ extend along the cell cortex to the cell posterior (Harris et al., 2016; Onishi et al., 2020). These EB1 comets likely track the ends of the dynamic cytoplasmic microtubules, rather than rootlets, because EB1 preferentially binds near the plus end of unstable, growing microtubules (Akhmanova & Steinmetz, 2008). Moreover, in dividing cells, EB1-NG colocalizes with the spindle and the cleavage furrow (Onishi et al., 2020), making it a marker to monitor mitotic events that may be controlled by CDKB1/CYCB1.

We used two different imaging methods to precisely record the behavior of EB1-NG in dividing wild-type and mutant cells. In method 1, we used 3-min intervals with single Z-planes to avoid phototoxicity, and the precise temperature control to image ts-lethal mutants through multiple division cycles, as described above in detail. Method 1 was also used for a movie with 20-sec intervals. In method 2, we used 10-sec intervals with single Z-planes to examine EB1-labeled structures that are near the medial plane (anterior spots, spindle, and furrow) in the first division of a multiple fission cycle (as described previously [Onishi et al., 2020]).

As cells enter mitosis, the polar ‘spot’ of EB1 signal splits into two; the two spots move slightly into the cell interior and mark foci that nucleate formation of a bipolar spindle about 4 min after pole splitting (Table 1, Figures 6 and 7, Supp Videos 3 and 4). Supp. Video 3 shows the process of pole splitting, spindle formation, anaphase and cytokinesis all marked by EB1-NG, at 10-sec time resolution in the first division cycle. Supp Video 4, at 3-min resolution, shows the same sequence repeating in three sequential divisions. (As noted in Methods, the higher time resolution resulted in sufficient irradiation of the cells that viability was lost; after the first division, additional divisions were rarely observed. Irradiating only every 3 min seemed to give division kinetics and numbers similar to those of unirradiated cells. Supp. Video 4 shows the high degree of synchrony of successive divisions, and the reliable appearance of the cytokinetic furrow at right angles to the dissociated spindle).

**Figure 6.**
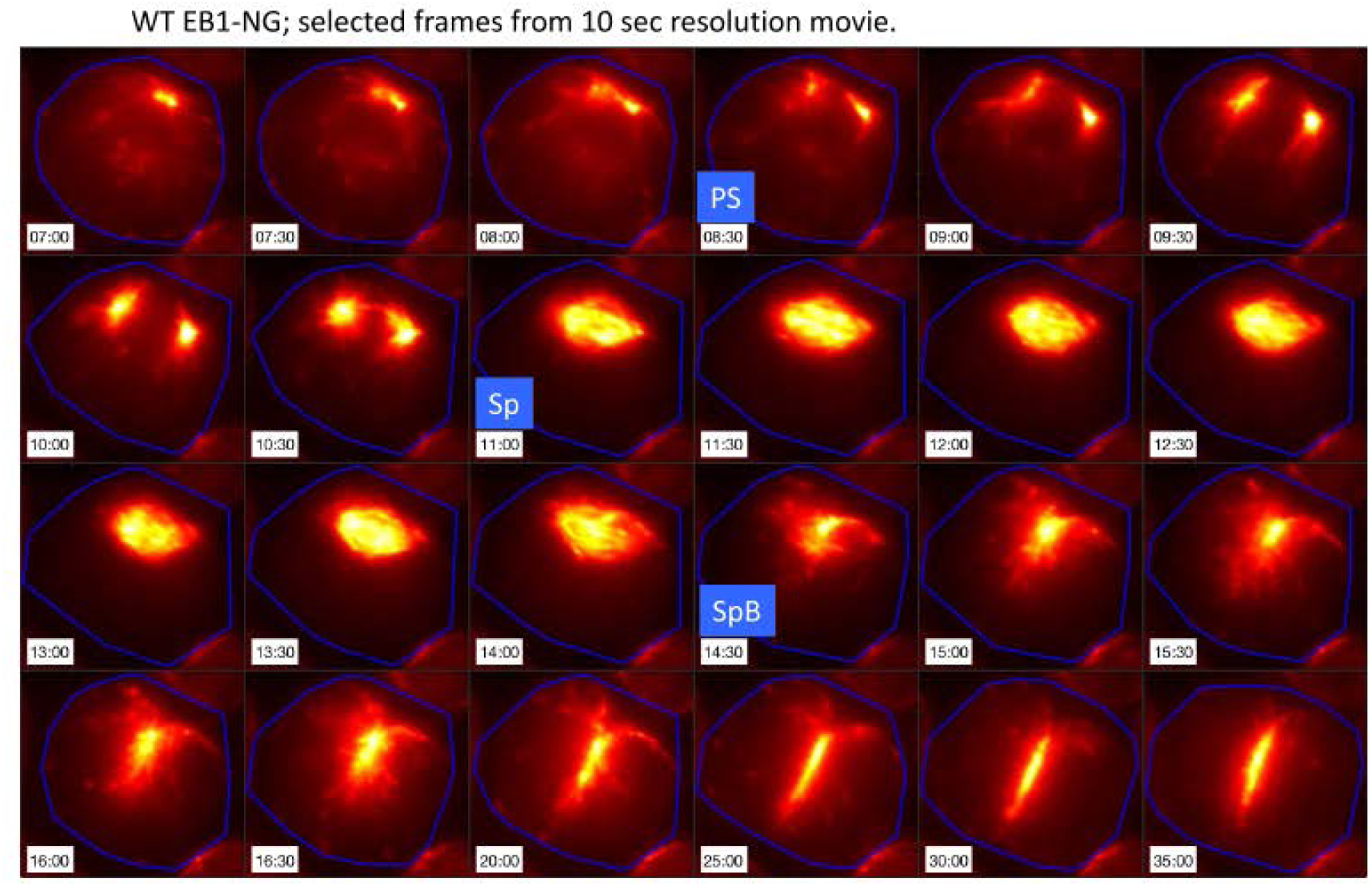
Live cell time lapse of wild-type EB1-NG cells with 10-sec. intervals. Live cell time lapse with 10-sec. intervals acquired with microscopy method 2 (see Methods section). EB1-NG signal in orange. PS: pole separation. Sp: spindle formation. SpB: spindle breakdown.

**Figure 7.**
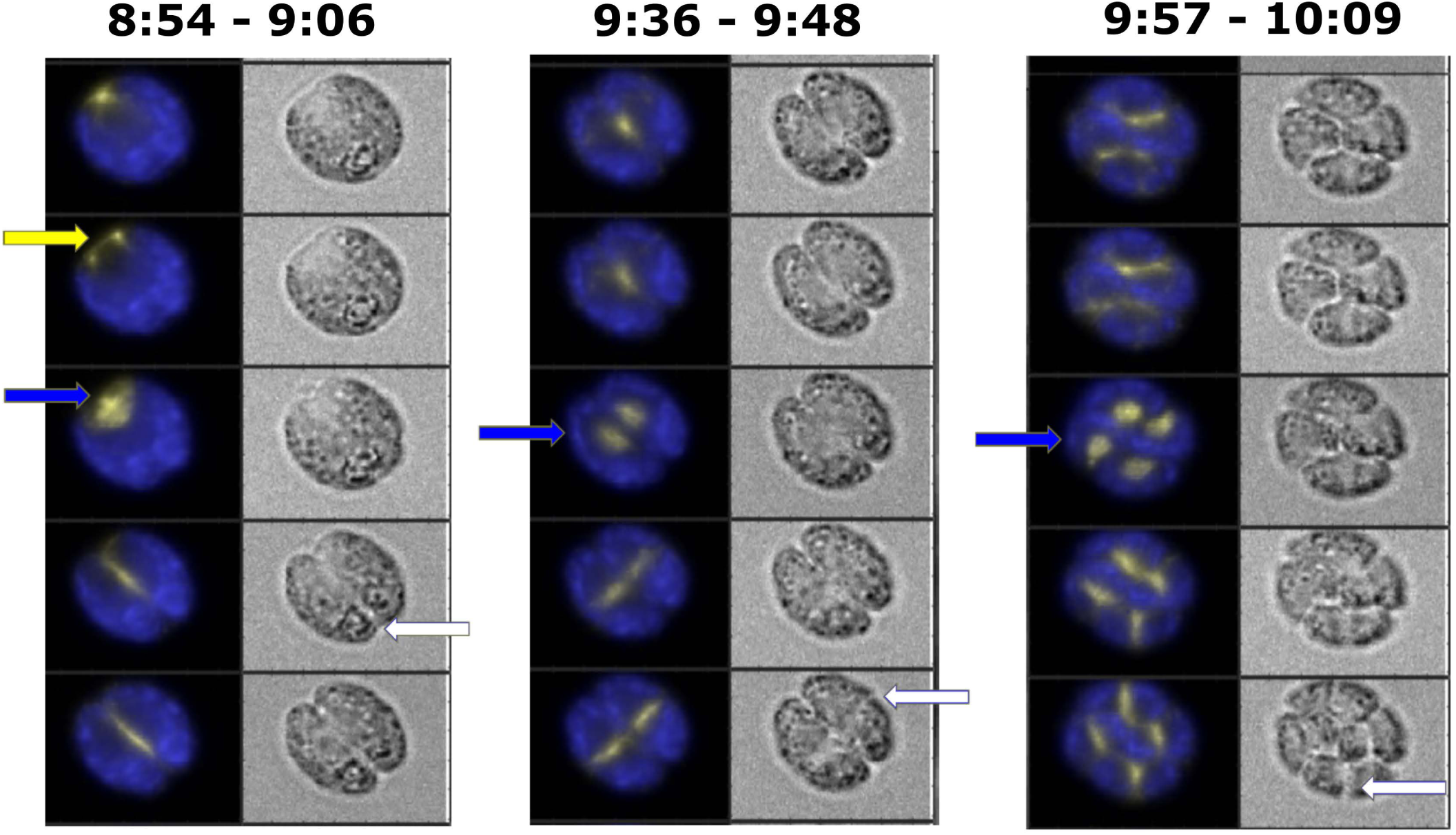
Live cell time lapse of wild-type EB1-NG cells with 3-min. intervals. Live cell time lapse with 3-min. intervals acquired with microscopy method 1 (see Methods section). Each cell has a brightfield image (right), and a composite of EB1-NG signal in yellow and chloroplast autofluorescence signal in blue (left). Yellow arrow: spindle pole separation. Blue arrows: new spindle formation. White arrows: new cleavage furrow formation. Time indicated on top of each strip is hours and minutes after beginning of time lapse. The indicated time corresponds to the top cell in each strip. Each subsequent cell going down is from an image captured every 3 minutes.

**Table 1.** Timing of EB1-scorable mitotic events. Time lapse movies at varying frame-rates (top row) were analyzed manually, and mean and standard deviation of time intervals is presented (all in minutes). ‘Pole sep’: separation of the anterior EB1 signal into two separate foci. ‘SP1, SP2, SP3’: full formation of bipolar spindle in 1^st^, 2^nd^, 3^rd^ rounds of division. A spindle was scored when the EB1-NG signal was continuous across the midline of the cell, and the orientation of the signal was roughly perpendicular to the following cleavage furrow. ‘SP1B, SP2B, SP3B’: spindle breakdown 1^st^, 2^nd^, 3^rd^ rounds of division. Spindle breakdown was scored when the EB1-NG signal was no longer perpendicular to the following cleavage furrow. Spindle formation and breakdown were highly synchronous in progeny within a single cell, although not all spindles were in focus in every division. To calculate the spindle duration, or SP to SPB, the frame number of the first visible spindle was subtracted from the frame number of the start of the spindle breakdown, and then multiplied by the frame frequency. In cases where a spindle was not visible in any frame it was given a value of 0 frames, but only if the preceding PS was clearly visible; in these cases, it was assumed that the spindle was likely missed because it was not present at the times the images were captured. If neither the PS nor the SP were seen, the values were not recorded. ‘CF1, CF2, CF3’: detectable cleavage furrow initiation in 1^st^, 2^nd^, 3^rd^ rounds of division (for later divisions, furrow formation in any progeny cell was counted due to image complexity). A CF was scored when a visible indent was observed in the cell membrane that was contiguous with the upcoming plane of separation of the cells. The 10-sec framerate movie was at 26°C; the 20-sec and 3-min movies were at 33°C. Entries are in minutes: mean +/- standard deviation (number of cells). ND: not determined, for the following reasons: (1) 10 and 20 sec framerates were only usable for the first division, as cell viability dropped from light exposure; (2) cleavage furrow formation was scored from brightfield images that were captured only at 3 min resolution. Effective detection of pole separation was difficult at 3 min resolution because it was easiest to observe when moving density could be compared between adjacent frames. Only the later phases of the PS were easily seen once the poles were separated and very bright. Therefore, this was not scored in the 3 min framerate movies. See Figures 6 and 7 and Supp. Videos 3 and 4 for illustrative examples.

| Event/Frame-rate | 10 sec | 20 sec | 3 min |
| --- | --- | --- | --- |
| Pole sep → SP1 | 3.3 (1) | 4 +/- 1 (14) | ND |
| SP1 → SP1B | 3.8 (1) | 3.7 +/- 0.7 (16) | 3 +/- 1 (30) |
| SP1B → CF1 | ND | ND | 1 +/- 1 (24) |
| SP1 → SP2 | ND | ND | 37 +/- 3 (18) |
| SP2 → SP2B | ND | ND | 2 +/- 1 (28) |
| SPB2 → CF2 | ND | ND | 2 +/- 2 (20) |
| SP2 → SP3 | ND | ND | 41 +/- 5 (15) |
| SP3 → SP3B | ND | ND | 3 +/- 1 (25) |
| SP3B → CF3 | ND | ND | 2 +/- 2 (23) |

The spindle structure has a ∼4 min lifetime, then disappears; signal remains at approximately the position of the spindle midzone, and this signal rapidly elongates perpendicular to the spindle axis (Figs. 6 and 7; Supp Videos 3 and 4). This line of EB1 signal is detected coincident with a cleavage furrow (detectable in a paired brightfield image), perpendicular to the former spindle axis. EB1 signal and the cleavage furrow extend essentially together in space and time (Fig. 7). This likely reflects growth of the microtubule array called the ‘phycoplast’, which marks (and is probably required for) cleavage furrow development (Ehler & Dutcher, 1998; Onishi et al., 2020). The four-membered rootlet microtubules run adjacent to the cytoplasmic microtubules in this array (Ehler et al., 1995), and may dictate its location (Ehler et al., 1995; Ehler & Dutcher, 1998).

Since formation of a cleavage furrow detectable in brightfield almost invariably occurs in the 3-min frame following spindle detection (Table 1), and CYCB1-GFP degradation tightly correlates with cleavage furrow formation (see above), these combined results suggest near-simultaneous spindle breakdown and CYCB1 degradation followed by cleavage furrow formation.

In multiple fission, additional cell division cycles occur within the same mother cell wall. These cycles are rapid and regularly spaced (Table 1). We almost invariably observe simultaneous appearance in the 2^nd^ division of two bipolar spindles, which disappear in the next frame replaced by two lines of EB1 signal perpendicular to the long axes of the spindles, and in most cells cleavage furrows were detectable in that same frame (Fig. 7, Supp. Video 4). In favorable 3^rd^-division cells, similar observations can be made of four bipolar spindles (Fig. 7). We expect that this reflects similar microtubule and EB1 behavior in the later divisions to what was observed in the first. However, at 3-min the frame resolution is close to the interval between pole splitting and spindle formation, and between spindle formation and breakdown; as a consequence in a sizable fraction of cells we observe pole splitting or spindle formation, but not both.

With a clear picture of the events and timing of EB1-labeled mitotic events in wild-type, we wanted to determine where various mutants were blocked in the progression. In *cycb1-5* cells expressing EB1-NG (Fig. 8, Supp. Video 5), we observed only the anterior spot of EB1-NG signal, as in WT G1 cells, even at long incubation periods (when WT cells have almost all divided). The polar splitting event was not observed.

**Figure 8.**
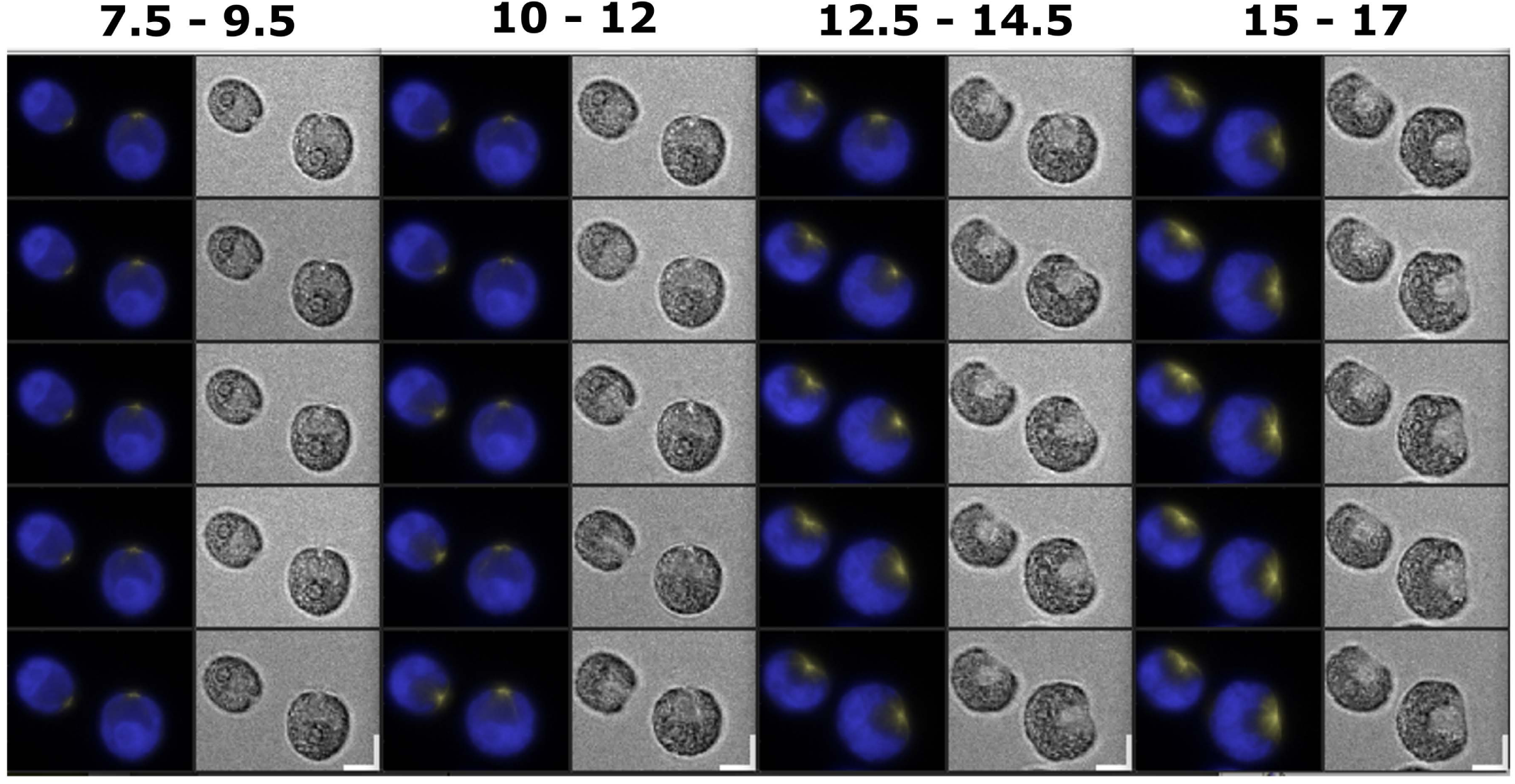
Live cell time lapse of cycb1-5 EB1-NG cells. Live cell time lapse acquired with microscopy method 1 (see Methods section). Each cell has a brightfield image (right), and a composite of EB1-NG signal in yellow and chloroplast autofluorescence signal in blue (left). Time indicated on top of each strip is hours after beginning of time lapse. The indicated time corresponds to the top cell in each strip. Each subsequent cell going down is from an image captured every 30 minutes.

To ask whether aspects of the *cycb1-5* phenotype might be due to failure to activate APC, we next turned to ts-lethal mutants of two genes required for APC function: *CDC27*, an essential core subunit of APC itself, and *CDC20,* an activator of APC. In both mutant backgrounds, cells expressing EB1-NG at restrictive temperature underwent the polar splitting reaction followed by efficient bipolar spindle formation (Figs. 9 and 10, Supp. Videos 6 and 7). Once formed, the spindle was stable, lasting for many hours (in contrast to ∼4 min in WT), consistent with previous observations with anti-tubulin immunofluorescence (Atkins & Cross, 2018). No cytokinetic cleavage furrow formed; correlated to this, there was no EB1-NG signal aligned perpendicular to the spindle long axis.

**Figure 9.**
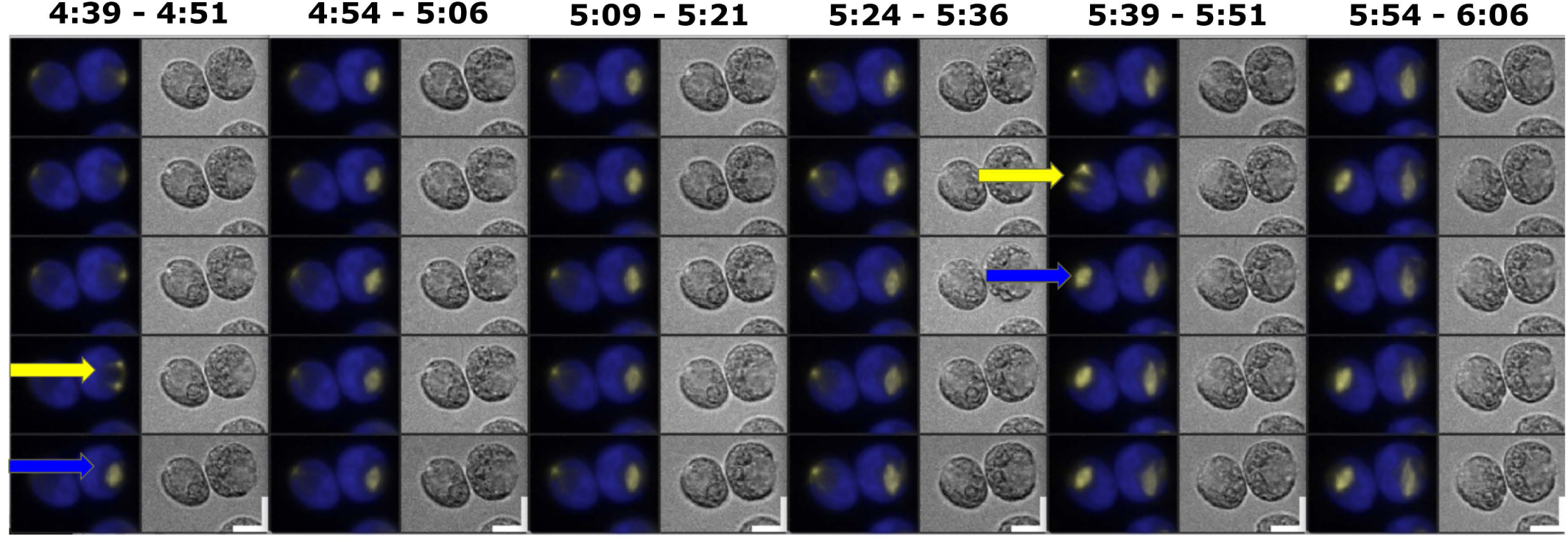
Live cell time lapse of cdc27-6 EB1-NG cells. Live cell time lapse acquired with microscopy method 1 (see Methods section). Each cell has a brightfield image (right), and a composite of EB1-NG signal in yellow and chloroplast autofluorescence signal in blue (left). Yellow arrows: spindle pole separation. Blue arrows: new spindle formation. Time indicated on top of each strip is hours and minutes after beginning of time lapse. The indicated time corresponds to the top cell in each strip. Each subsequent cell going down is from an image captured every 3 minutes.

**Figure 10.**
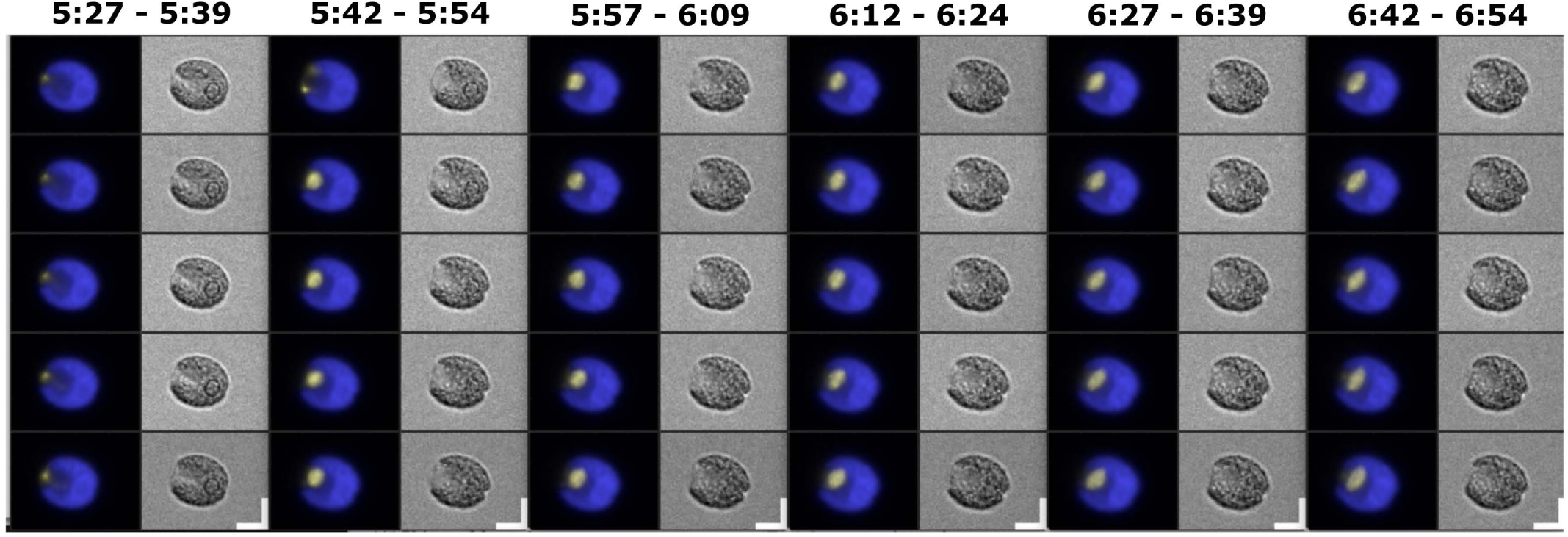
Live cell time lapse of cdc20-1 EB1-NG cells. Live cell time lapse acquired with microscopy method 1 (see Methods section). Each cell has a brightfield image (right), and a composite of EB1-NG signal in yellow and chloroplast autofluorescence signal in blue (left). Time indicated on top of each strip is hours and minutes after beginning of time lapse. The indicated time corresponds to the top cell in each strip. Each subsequent cell going down is from an image captured every 3 minutes.

*Chlamydomonas* has a gene orthologous to the *CDC20* homolog *CDH1*, which in other organisms can also activate the APC. The largely consistent results with EB1 comparing *cdc20-1* and *cdc27-6* mutants suggest that these phenotypes are due to loss of the APC^CDC20^ complex, and that *Chlamydomonas* CDH1 is unable to fully substitute for CDC20. This is also true in yeast and animals, owing at least in part to CDK-dependent inhibition of CDH1-APC (Zachariae & Nasmyth, 1999).

In interphase, EB1-NG is associated with ‘comets’ that move from the anterior along the cortex to the posterior (Harris et al., 2016), presumably marking rapid microtubule growth. These comets disappear for a very brief interval exactly coincident with presence of a spindle (Fig. 11A, C; Supp. Fig. 3). In *cdc20-1* cells the spindle is stable, and comet suppression is permanent (Fig. 11B, D). This suggests APC-Cdc20-dependent degradation of an inhibitor of comet formation.

**Figure 11.**
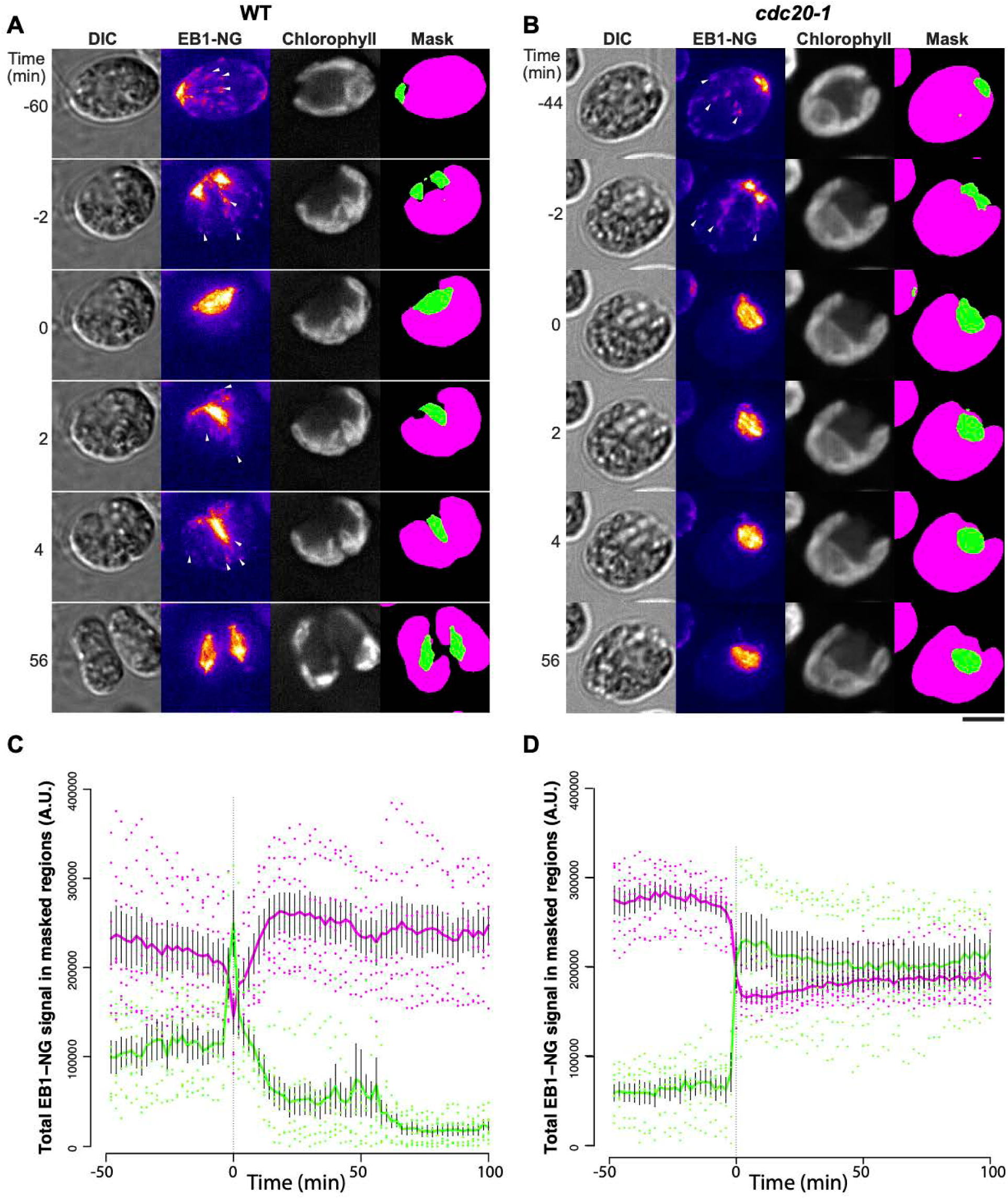
Suppression of cytoplasmic EB1-NG comets during mitosis. (A, B) Representative examples of WT (A) and cdc20-1 (B) cells expressing EB1-NG. Time-lapse microscopy was done using Method 3 (see Materials and Methods). Bar, 5 µm. (C, D) Regions representing polar dot, spindle, and furrow (green in A, B) and cytoplasm (magenta in A, B) were masked as described in Materials and Methods. Total signals in the masked regions are presented as mean ± SEM (N = 7), with values from individual cells overlaid as dots. Time zero (first appearance of complete spindle) was determined empirically for each cell. See Supp. Figure 3 for individual traces.

### Simultaneous localization of CYCB1 and EB1

To examine if there is colocalization between CYCB1 and EB1, we constructed a CYCB1-GFP EB1- mScarlet strain and observed the first division with live cell microscopy (microscopy method #3 with z-stacks). The localization of EB1-mScarlet is similar to what was observed above with EB1-NG: a polar signal splits into two as cells enter mitosis, and a bipolar spindle is formed from these two spots; the spindle then disappears and EB1 collects along the cleavage furrow (Figs. 6, 12; Onishi et al. 2020).

**Figure 12.**
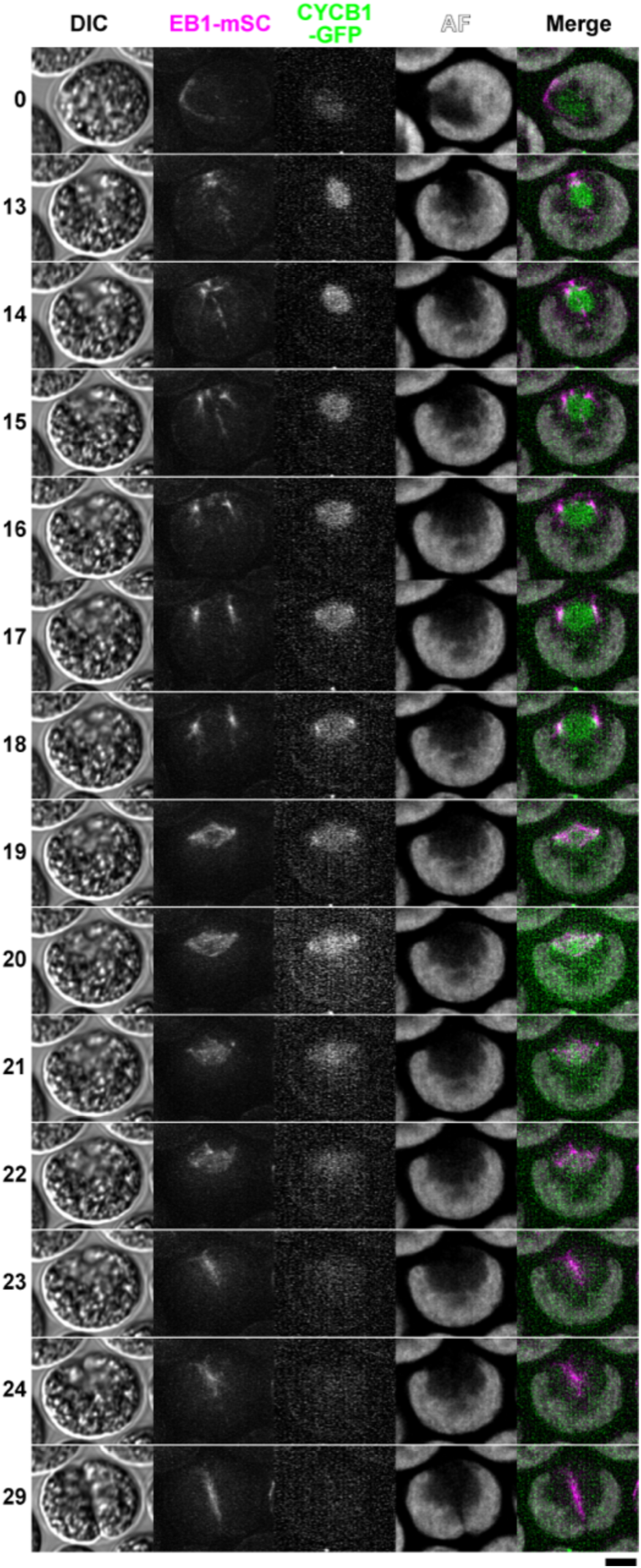
Live cell time lapse of CYCB1-GFP EB1-mScarlet cells with 1-min intervals. Live cell time lapse imaging was done using Method 3 (see Methods section). Imaging was done at 27 C. Select time-frames are shown with the times indicated on the left (min.). For DIC and AF (chlorophyll autofluorescence), mid-section images are shown; for EB1-mSc, maximum projections of Z-stacks are shown; for CYCB1-GFP, maximum projections of Z-stacks after Gaussian blurring along the Z-axis are shown. Bar, 5 microns. See Supp. Video 8 for a larger field of view including this cell and the entire time-series.

CYCB1-GFP signal is briefly concentrated at or just adjacent to the EB1 foci, just before spindle formation (Fig. 12, timepoints 18-20 min; Supp. Video 8). This places CYCB1-GFP at or near the future spindle poles. As the spindle breaks down, CYCB1-GFP shows transient localization to a band at the approximate location of the former spindle midzone; CYCB1-GFP is then completely degraded. Thus CYCB1 degradation and spindle breakdown are nearly simultaneous, in agreement with our conclusions comparing separate CYCB1-GFP and EB1-NG movies (see above).

The genetic requirement for CYCB1 for spindle formation, and the localization of CYCB1 to spindle poles just before spindle formation, suggests the speculation of direct regulation of spindle assembly by pole-localized CYCB1.

## DISCUSSION

### CYCB1 interacts with CDKB1

Previously (Atkins & Cross, 2018), we inferred that CYCB1 was the most likely activator of CDKB1 enzymatic activity, since CDKB1-associated histone H1 kinase activity was greatly reduced in immunoprecipitates from *cycb1-5* cells. Here, we confirm and extend this finding: CYCB1 binds CDKB1 but not CDKA1 in immunoprecipitates from doubly tagged cells, and CYCB1-associated histone H1 kinase activity is absent in immunoprecipitates from *cdkb1-1* cells.

### Cell-cycle-regulated and APC-dependent CYCB1 proteolysis

APC inactivation greatly increased CDKB1-associated kinase activity, without altering CDKB1 levels (Atkins & Cross, 2018), and we inferred that this was likely due to blocking APC-dependent CYCB1 proteolysis. Here we show that CYCB1-GFP abundance is indeed sharply cell-cycle-regulated. For a brief period surrounding cell division, the estimated half-life of CYCB1-GFP is reduced to ∼3-5 min., and degradation is absolutely dependent on functional APC. Turnover in each cell cycle cannot be detected in bulk culture (Fig. 1) because synchrony is insufficient. This is to be expected since the period of instability is unlikely to be longer than 10-20% of each division cycle, so even slight offsets in timing among cells will prevent detection of synchronous degradation. Notably, though, CYCB1-GFP degradation is sharply synchronous within daughter progeny in a single cell undergoing multiple fission. The basis for synchrony (whether due to identical timing in independent progeny cells, or communication between cells) is unknown.

Classic experiments in *Xenopus* embryos show that cyclin B is needed for mitotic entry, but its degradation is needed for mitotic exit (Murray et al., 1989; Murray & Kirschner, 1989). In budding yeast, the degradation of the B-type cyclin Clb2 is essential for viability (Wäsch & Cross, 2002), though this requirement can be bypassed by periodic inhibition of cyclin B-CDK-associated kinase (Thornton & Toczyski, 2003). The closely-related B-type cyclin Clb3, however, can persist undegraded without blocking mitotic exit, and without impacting viability (Pecani & Cross, 2016). In *N. tabacum*, expression of destruction-box-deleted *CYCB1* resulted in defective cytokinesis (Weingartner et al., 2004), perhaps due to inactivation of cytokinesis-inducing proteins by CDK phosphorylation (Sasabe & Machida, 2014). A requirement for cyclin B degradation for cytokinesis could account for an essential requirement for degradation. Independently, relicensing of replication origins is also blocked by high CDK activity in many systems (Kearsey & Cotterill, 2003), providing a distinct reason why cyclin B degradation might be essential.

Inability to complement *cycb1-5* ts-lethality by transformation with *CYCB1-db-GFP* (with the conserved destruction box deleted) is consistent with CYCB1 degradation being essential in *Chlamydomonas* as well. We speculated previously that CYCB1 might inhibit completion of cytokinesis, since elevating CYCB1 levels by *apc* inactivation is associated with absence of a cleavage furrow, while *apc cycb1-5* double mutants form an aberrant partial furrow similar to that produced by *cycb1-5* single mutants (Atkins & Cross, 2018).

### CDKB1 levels are regulated by entry into ‘division phase’ but not by cell cycle position

*CDKB* transcription and protein accumulation is elevated in mitotic cells in *Cyanodioschizon*, *Ostreococcus, Physcomitrella,* and *Arabidopsis* (red and green algae, moss, and land plant) (Corellou et al., 2005; Nowack et al., 2012). This leads to the model that its degradation after mitosis could be cell-cycle-phase-specific, perhaps serving the same function as cyclin B degradation, to allow mitotic exit (Adachi et al., 2006; Corellou et al., 2005). Our results show that in *Chlamydomonas,* this is not so. Unlike cyclin B, CDKB1 (the sole CDKB family member) is not removed at the conclusion of each mitosis. Rather, CDKB1 is restricted to what we call ‘division phase’: a condition of commitment to cell divisions (whether one or many) (Cross, 2020; Heldt et al., 2020). Cells undergoing multiple divisions make CDKB1 before the first division, and it stays high until all divisions are complete (and cells exit ‘division phase’); it is then rapidly degraded. The distinction between mitosis-specific accumulation and division-phase-specific accumulation is more easily made in *Chlamydomonas* as a result of multiple fission biology.

We recently suggested an equivalence between classical ‘commitment’ to division, and activation of transcription of a large number of division-essential genes including *CDKB1* (Cross, 2020). We speculate that transcription of these genes may be continuous throughout the period of multiple fission; lack of any drop in CDKB1 protein levels between divisions is consistent with this idea. We found recently that the replication control protein MCM4, a member of the mitotic transcriptional regulon along with CDKB1, accumulates as cells enter division phase, remains at a high level until the terminal division, then is degraded (Ikui et al., 2021), thus exhibiting similar behavior to CDKB1. This is consistent with the idea of ‘division phase’ as a discrete cellular state, permissive for cell cycle progression but independent of specific cell cycle phase (Cross, 2020; Heldt et al., 2020). CDKB1 lacks a recognizable target for APC-dependent degradation (D-box or KEN box) but nevertheless it is not degraded in a *cdc27-6* background (Figs. 1, 4). Thus, there must be a separate pathway between APC and CDKB1. The mitotic transcriptome continues to be expressed at a high level in this background (Tulin and Cross 2015; FC and KP, unpublished results); thus, the APC is required for exit from division phase, and CDKB1 may remain stably accumulated at a high level for this reason.

### Regulation of microtubule dynamics and morphogenesis by CYCB1-CDKB1 and APC-CDC20

EB1-NG was shown to be an informative single-cell marker for mitotic progression in *Chlamydomonas* (Onishi et al., 2020). As cells entered mitosis EB1-NG localization undergoes dramatic changes (Onishi et al., 2020): the single polar focus of EB1-NG splits into two and separates; the mitotic spindle then forms between these two foci, and persists for ∼4 min before anaphase. Specifically during this period, the cortical comets characteristic of interphase cells (Harris et al., 2016) are entirely suppressed (Fig. 11). After spindle breakdown, EB1-NG signal immediately moved to a line perpendicular to the former spindle axis, and marked the growing cleavage furrow. Cleavage in *Chlamydomonas* is strongly dependent on microtubules (Ehler & Dutcher, 1998), and can occur in the complete absence of F-actin (Onishi et al., 2020), so the furrow localization of EB1-NG likely reflects essential microtubule growth during cytokinesis.

CYCB1/CDKB1 is required for the first step in this process; arrested *cycb1* cells keep a single anterior focus which does not split (Fig. 8). Because no spindle forms in these cells, polar splitting in wild-type may produce poles required for spindle generation. The mutant cells form an initial cellular indentation (the ‘notch’; Tulin & Cross, 2014) (Fig. 8), at the position of the anterior EB1-NG focus, but no extension of a line of EB1-NG into the cell (as observed in full cytokinesis; Onishi et al.,2020; Fig. 7) is observed.

Localization of CYCB1 to the region of the spindle poles 1-2 min before spindle formation is consistent with a direct regulation of the microtubule-organizing activity of spindle poles. In animal cells and in yeast, cyclin B localizes to centrosomes via the cyclin B ‘hydrophobic patch’ docking motif (Basu et al. 2020 and references therein); and *Chlamydomonas* CYCB1 retains all key residues making up the hydrophobic patch. In contrast to *Chlamydomonas*, though, this localization may occur long before actual spindle formation. It is also important to note that in *Chlamydomonas*, the spindle pole is spatially distinct from the basal body (centrosome equivalent) (O’Toole and Dutcher 2014). We do not know whether CYCB1 localization is specific to one or the other of basal bodies or spindle poles.

Inactivation of APC or CDC20 has no effect on EB1 polar splitting or spindle formation, but anaphase, cleavage and cytokinesis are completely blocked; consistently, no ‘line’ of EB1-NG signal perpendicular to the spindle axis is observed in these blocked cells (Figures 9 and 10).

These results imply strong and opposing effects on microtubule dynamics and morphogenesis by CYCB1/CDKB1 versus APC-CDC20. These events occur in stereotyped time intervals coordinated (and likely caused by) tight sequential changes in CYCB1 levels and APC-CDC20 activity.

Regulation of microtubule dynamics and spindle morphogenesis by cyclin B-CDK may be conserved throughout eukaryotes (Basu et al., 2020; Verde et al., 1992) – our results extend this conservation to the deeply diverged plant kingdom. Overall, we observe strong conservation between *Chlamydomonas* and yeast and animals of the roles of cyclin B/CDK and the APC with respect to their inter-regulation and their overall effects on cell cycle biology including spindle morphogenesis. However, while cyclin B has a highly conserved role in mitosis, its associated kinase subunit is CDKB1 rather than CDK1/CDKA1 as in yeast and animals; this substitution may be universal within the Viridiplantae plant kingdom (Atkins & Cross, 2018; Corellou et al., 2005; Nowack et al., 2012; Tulin & Cross, 2014). In the plant kingdom, CDKA1 may instead be specific to cell size control and the G1/S transition (Cross, 2020). *Chlamydomonas* thus provides a unique opportunity to investigate the molecular regulation and mitotic functions of the plant-specific mitotic inducer CYCB/CDKB in a unicellular system, at high spatial and temporal resolution.

## Materials and Methods

Immunoblotting, immunoprecipitation, and protein kinase assays were carried out as previously described (Atkins & Cross, 2018).

### Fluorescent reporter constructs

#### CYCB1-GFP

We constructed a plasmid with 1.3 kb of genomic DNA upstream of *CYCB1*, followed by the *CYCB1* coding sequence with introns; the termination codon was replaced with 3 copies of a GlyGlyGlyGlySer linker sequence followed by GFP. After the GFP termination codon the plasmid contained 1.1 kb of the 3’ UT region from *CDKB1*, followed by a 1 kb fragment containing a paromomycin resistance cassette.

We linearized this plasmid and transformed a *cycb1-5* strain by electroporation as described (Atkins & Cross, 2018). We recovered transformants in two ways: either by selection on paromomycin at 21 degrees (permissive temperature for *cycb1-5*) or by selection without paromomycin at 33 degrees (non-permissive temperature). For unknown reasons, likely related to the known fragmentation of transforming DNA in *Chlamydomonas*, all of the paromomycin-resistant colonies tested were temperature-sensitive, and none of the temperature-resistant colonies were paromomycin-resistant. We chose one temperature-resistant transformant and found linkage in tetrad analysis between a single locus containing *GFP* by PCR, and rescue of temperature-sensitivity of *cycb1-5*. Parallel transformations with an identical plasmid with a deletion of the *CYCB1* destruction box (Atkins and Cross, 2018) failed to yield any temperature-resistant transformants in multiple experiments.

*Chlamydomonas* transgenes are frequently subject to random silencing (Schroda, 2019). We largely eliminated this problem with *CYCB1-GFP* by selection of cultures at non-permissive temperature before time-lapse microscopy (see below). Even with this precaution, we observed sporadic cells in time-lapse that failed to express *CYCB1-GFP,* instead arresting with the characteristic morphology of *cycb1-5* (Atkins & Cross, 2018).

In one experiment in this paper (Fig. 12, Supp. Video 8) we used an allele of *CYCB1-GFP* in which the endogenous copy of *CYCB1* was tagged with GFP, by a method to be described elsewhere (MO and FC, unpublished). The endogenously tagged *CYCB1-GFP* behaved similarly to the *CYCB1-GFP* transgene used in all other experiments reported here.

#### EB1

To construct pMO699 (EB1-mSC), the mNeonGreen sequence in pCrEB1-NG (Harris et al., 2015) was excised out using *Xho*I sites and replaced with mScarlet-I [amplified from mScarlet-I-mTurquoise2 (Addgene, Plasmid #98839) (Mastop et al., 2017] by Gibson assembly. pMO669 was then linearized using *Eco*RI and *Sca*I prior to transformation into *Chlamydomonas* by electroporation.

In the experiment shown in Fig. 11, an endogenously tagged *EB1- NG* allele was employed, constructed by a method to be described elsewhere (MO and FC, unpublished). As with *CYCB1-GFP*,the endogenously tagged protein behaved similarly to transgene used in all other experiments reported here.

### Time-lapse Microscopy

Multiple imaging methods were used. Method #1 was used for single Z-plane imaging at 3 min. intervals and low fluorescence exposure times to avoid cell phototoxicity and to image cells through multiple division cycles. Method #2 was used for 10- or 20-second interval movies at a single Z-plane. Method #3 was used for 1-minute interval movies with high fluorescence exposure times. These methods are complementary; frequent exposures, high exposure times and multiple Z planes allowed high-resolution detection of events within a single cell division, but the imaged cells generally lost viability soon after; while the much lower overall illumination of Method #1 reduced temporal and spatial resolution but allowed reliable imaging of an entire multiple fission cycle (at the end of which viable cells hatched from the mother and swam away).

**Method #1:** Cells were taken from a 2-day culture on a TAP plate, transferred to liquid TAP for 4 hrs. for cells to become motile, then swimming cells were separated from dividing and other non-motile cells and debris. This separation was achieved by pipetting 500 µL of the liquid cell culture, removing the pipette tip from the pipette, then placing the pipette tip into another tip containing 500 µL TAP + 2% Ficoll, such that the end of the pipette tip containing the cells was in contact with the Ficoll. A white LED (Evan Designs) was then placed on the wide end of the pipette tip pair. This complete apparatus was then put in a dark enclosure. With the LED being the only source of light inside the enclosure, cells swim away from the light, into the Ficoll, and collect on the end of the pipette tip. Once a sufficient number of motile cells had traversed the Ficoll and accumulated on the pipette tip, the pipette tip was removed, and the cells were pushed out by lightly pressing the wide end of the pipette tip. This ‘swim-selected’ population mostly consisted of small to medium motile cells (due to the high density of the Ficoll), with very few large or dividing cells carried over by the flow of the smaller cells.

To immobilize the swim-selected cells for long-term time lapse microscopy, they were placed on agarose medium immediately after collection, similar to what was used by Di Talia et. al. (2007) for budding yeast microscopy. However, the setup employed by Di Talia et. al. (2007) involved large agarose slabs placed close together on a glass cover slip, and covered with a clear plastic piece, which was then sealed along the edges with paraffin. This setup could not be used for long-term microscopy of *Chlamydomonas* for the following reasons. First, unlike budding yeast, *Chlamydomonas* cells are motile, so placing agarose slabs close together on a cover slip allows for the possibility of cells swimming from one slab to the other (if a connective layer of liquid is formed between the slabs). Second, in order to make long-term movies (20 hrs.), drying of the agarose must be minimized. A large plastic cover placed over the agarose slabs allows for enough drying during the course of the movie that cells often drift completely out of the field. Drying causes cells to move along the z-axis as well, necessitating a very large autofocus range. A large plastic cover also collects water on its inner surface by condensation, which was significant at the temperature at which we intended to make movies (33.3°C).

To avoid these issues, we designed a small cylindrical chamber with one open side (inner diameter: 5mm; inner height: 3.5 mm; wall diameter: 1 mm). The chamber was fabricated from clear acrylic sheets using a laser cutter at Rockefeller University’s Precision Instrumentation Technologies facility. The barrel portion of the chamber was made by cutting two concentric circles on a 3.5 mm-thick acrylic sheet. The inner circle had a diameter 2 mm smaller than the outer circle, so that the sides of the barrel were 1 mm wide. A lid was made by cutting a clear 1 mm-thick acrylic sheet in a circle with a diameter equal to the outer diameter of the barrel. The lid was then attached to the barrel with acrylic cement (Scigrip).

Molten TAP with 1.5% SeaKem NuSieve GTG agarose (Lonza) was poured into the chamber to a level of 3 mm above the top and a glass slide was placed 1.5 mm above the upper edge of the box, flattening the agarose. The agarose was allowed to solidify at room temperature for 10 min., then the glass slide was removed. Cells were pipetted (0.5 µL) onto the agarose surface and kept at room temperature for 15 min. to allow the surface to dry. The agarose edges were trimmed so that the exposed agarose surface was flat throughout. The cell side of the box was placed onto a 24 x 50 mm glass cover slip (Fisherbrand) and the exposed agarose portion was sealed with VALAP (equal mass petroleum jelly, paraffin, lanolin). When multiple cell chambers were used, they were placed 1 cm apart (center-to-center). Plastic cover slips (Rinzle and ACLAR, both from Electron Microscopy Sciences) occasionally resulted in better cell viability and division number compared to glass (mostly 3 divisions compared to 2 divisions on glass), but this difference was irregular between batches. Glass cover slips from our current supplier (Fisher Scientific ‘Fisherbrand’ Cat. No. 12-545-F), have been consistently better than Rinzle or ACLAR plastic in maintaining cell viability and most cells divide 3-4 times. Glass has the additional benefits of lower autofluorescence compared to plastic, and less flexion, which results in less drift along the z-axis, making autofocusing easier.

Time lapse microscopy was carried out on a Leica DMI6000B inverted microscope, using a 63X objective, with the objective and stage heated to 33.3°C. Images were acquired using custom software, as previously described for budding yeast microscopy (Charvin et al., 2008), but with modifications to improve autofocus for *Chlamydomonas*. We acquired brightfield images instead of phase contrast, because brightfield allowed for more reliable autofocusing. Fluorescence images were acquired using a Leica EL6000 mercury-arc lamp and a 30% neutral density filter. GFP images were acquired with 0.4 s exposure using a narrow-band eGFP filter set from Chroma (Cat. No. 49020) to minimize autofluorescence. Venus and mNeonGreen (NG) images were acquired with 0.3 s exposure using an eYFP filter set from Chroma (Cat. No. 49003). For chloroplast background, we acquired images with 0.003 s exposure using a Cy5 filter set from Chroma (Cat. No. 49006).

In *Chlamydomonas*, chloroplasts fluorescence is detectable at most wavelengths, and this seriously interferes with detection of the rather dim CYCB1-GFP signal. We developed a simple deconvolution procedure to subtract chloroplast background from GFP (see below).

To provide the cells with illumination for photosynthesis between frames, we placed white LEDs (Evan Designs) 10 mm above the cell chambers and 7 mm away from the imaging axis, so that the irradiance at the location of the cells was 150 µmol photons m^-2^ s^-1^. The LEDs were mounted on a 3D-printed plastic enclosure that covered the cell chamber. The transmitted light path from above was not impeded because a clear plastic ACLAR film (Electron Microscopy Sciences) was used as a top. This enclosure also helped maintain temperature stability of the cell chamber by partially insulating against ambient temperature fluctuations. The LEDs were connected to a computer-controlled on/off timer (PowerUSB). The LED lights were off for the duration of the transmitted light/fluorescence image acquisition, then on for most of the remaining time until the subsequent frame. Because of imperfect synchrony between the time lapse image acquisition schedule and the exterior LED light on/off timer, 10-20 sec. were added to the LED off time, allowing a minimum of 90 sec. of LED illumination between 3-min. frames.

Temperature stability and accuracy during the course of a time lapse movie was extremely important. We found that in the microscopy setup described above, wild-type cells are inviable at 34°C and above. Many temperature-sensitive mutants do not arrest tightly below 33°C. Therefore, our movies were done at 33.3°C (± 0.3°C). To measure the temperature exactly at the location of the cells, we embedded a 0.1 mm diameter thermocouple (PerfectPrime TL0201) in the agarose microscopy chamber. To maintain this small temperature range, we heated the objective (with an aluminum collar) and stage (with an aluminum insert) with Peltier modules run by Oven Industries 5C7-195 controllers. To minimize the effect of air currents above and below the stage, we covered openings below the stage with aluminum foil, and used a 3D-printed plastic enclosure above the stage. The enclosure was printed at Rockefeller University’s Precision Instrumentation Technologies facility.

**Method #2:** As described previously (Onishi et al., 2020).

**Method #3:** Cells were synchronized using the 12L:12D light cycle at 26°C. At ∼11 h, the cells were collected by centrifugation and spotted on a small block of TAP + 1.5% low-melting-point agarose (Bio-Rad), which was then placed in a glass-bottomed 18-well chamber (Ibidi) and sealed with additional TAP + low-melting-point agarose. Imaging was done using a Leica Thunder inverted microscope equipped with an HC PL APO 63X/1.40 N.A. oil-immersion objective lens and an OkoLab incubator chamber that was maintained at 27°C. Signals were captured using following combinations of LED excitation and emission filters: 510 nm and 535/15 nm for CYCB1-GFP and EB1-NG; 550 nm and 595/40 nm for EB1-mSc; 640 nm and 705/72 nm for chlorophyll autofluorescence (AF). Time-lapse images were captured at 2-min intervals with 0.6 µm Z-spacing covering 9 µm; still images were captured with 0.21 µm Z-spacing covering 10-15 µm. The acquired fluorescence images were processed through Thunder Large Volume Computational Clearing and Deconvolution (Leica). Background chloroplast signal was removed from GFP images essentially as described below. Maximum projections from 15 z-stacked images of CYCB1-GFP and EB1-mSC were used in Fig. 12 and Supp. Video 8. CYCB1-GFP maximum projections were grainy because the signal was close to background; this problem was reduced by a Gaussian blurring of the GFP stack before the maximum projection was calculated (0.5 * image (n-1) + image(n) + 0.5 * image(n)).

#### Quantification of EB1-NG signals

The “Peak” mask representing the polar dots, mitotic spindle, and furrow, was created from MAX-projected EB1-NG images by applying Gaussian blur filtering (1.5 pixels) and a Default thresholding filter in ImageJ. A mask representing the total cell body was generated from MAX-projected AF images by applying Gaussian blur filtering (2 pixels) and a Triangle thresholding filter. Subtraction of the “Peak” region from this mask yielded a mask essentially covering the cytoplasm. Unlike the mid-section images shown in the figures, this MAX projection covers most of the cell body after binarization using an appropriate threshold, except that the thin cortical layer is not covered. Strong 2-pixel blur was applied to expand the signal so that the resulting mask covers the cortex. In interphase cells, EB1 signal under this cytoplasmic mask was mainly due to discrete ‘comets’ of EB1 traveling along the cortex to the cell posterior (Harris et al., 2016). Signals of EB1-NG were quantitated in each mask after uniform subtraction of background corresponding to intensities in non-cell areas.

### Deconvolution for Time Lapse Image Analysis

Autofluorescence from chloroplasts accounted for a large majority of the total signal with CYCB1-GFP or CDKB1-Venus detection. We developed a simple computational deconvolution procedure that largely corrected this problem. The key observation is that due to the broad excitation and emission spectra of photosynthetic pigments, chloroplasts are detectable with filters specific for GFP, YFP or RFP; in contrast, GFP and YFP have no signal under RFP detection. The brightest RFP signal was invariably detected in the posterior region of the cell where chloroplasts are known to reside. Therefore, assuming that chloroplast pigments have the same ratio of GFP:RFP detection at all points in the cell, it is straightforward, given paired images for GFP and RFP detection, to determine this ratio from high-RFP pixels (presumably deriving purely from chloroplast), and then to deconvolve the GFP-specific signal throughout the image (Supp. Fig. 2). This deconvolution is carried out automatically using the same algorithm for every image. To account for possible variations in lamp intensity or exposure time through a movie, the deconvolution ratio is calculated separately for each image in the series. Suppose F is the average ratio of red to green signal in the pixels with the highest red signal (pure chloroplast). Consider another pixel potentially containing both chloroplast and CYCB1-GFP signal. If CYC is the amount of CYCB1-GFP contributing to signal from that pixel, and the total green and red signals from that pixel are G and R respectively, then

G = FR + k*CYC, where k is a constant reflecting green emission from a given amount of CYCB1-GFP. Therefore, amount of CYCB1 in that pixel (in arbitrary units) is:

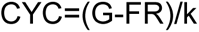

Assuming similar lamp intensity and exposure through the movie, k is a constant that is buried in arbitrary units for CYCB1-GFP. Given the assumptions above, this results in a linear measure of CYCB1-GFP comparable across an image and between images in a series, with the contribution of chloroplast to green signal removed.

The same procedure works for YFP.

MATLAB code to carry out the deconvolution is available on request.

The authors responsible for distribution of materials integral to the findings presented in this article in accordance with the policy described in the Instructions for Authors (https://academic.oup.com/plcell/pages/General-Instructions) are: Fred Cross; Masayuki Onishi.

## Acknowledgments

We thank Karl Lechtreck for sharing pEB1-NG and an anti-EB1 antibody. We also thank the Chlamydomonas Resource Center for providing essential strains and reagents. This work was supported by National Science Foundation Grant MCB 1818383 (to M.O. and John R. Pringle), by Duke University Department of Biology, and by PHS grant RO1GM078153 to FC.

## FIGURE LEGENDS

**Supp. Figure 1. Live cell time lapse of CYCB1-GFP in a *cdkb1-1* background**

Live cell time lapse acquired with microscopy method 1 (see Methods section). Each cell has a brightfield image (right), and a composite of EB1-NG signal in yellow and chloroplast autofluorescence signal in blue (left). Time indicated on top of each strip is hours and minutes after beginning of time lapse. The indicated time corresponds to the top cell in each strip. Each subsequent cell going down is from an image captured every 3 minutes.

**Supp. Figure 2. Illustration of subtraction of chloroplast autofluorescence in microscopy images**

Autofluorescence subtraction method demonstrated using single CYCB1-GFP cell. First column: RFP detection (colored blue) (chloroplast signal only). Second column: GFP detection channel only (CYCB1-GFP signal + chloroplast signal). Third column: composite of first two columns. Fourth column: GFP signal remaining after deconvolution (removal of contribution of chloroplast signal to GFP channel, leaving CYCB1-GFP signal only). Fifth column: composite of RFP channel (chloroplast only) and deconvoluted green channel (CYCB1-GFP signal only). Bottom: quantification of total RFP and GFP signals (left), and GFP signal concentration (blue line) before and after deconvolution.

**Supp. Figure 3. Traces for individual cells used for quantification in Figure 11.**

Top and bottom rows show the total signal under the green and magenta masks in Figure 11, respectively.

**Supp Video 1. CYCB1-GFP through 3 divisions.** Top left: blue line: RFP (chloroplast) signal. Yellow line: deconvolved CYCB-GFP signal. Top 2^nd^ graph: yellow: total GFP signal; black: a minimal convex hull was calculated containing 50% of the total cell GFP signal and concentration calculated; 3^rd^: histogram of intensities in the convex hull; 4^th^: surface plot of GFP intensity. Below: brightfield (left) and fluorescence (right). Black line: manually selected cell outline. White line: computed convex hull.

**Supp Video 2. CYCB1-GFP in *cdkb1-1* background.** Graphs and images as in Supp. Video 1.

**Supp. Video 3. Live cell time lapse of wild-type EB1-NG cells with 10-sec. intervals**

Live cell time lapse with 10-sec. intervals acquired with microscopy method 2 (see Methods section). EB1-NG signal in blue.

**Supp. Video 4. Live cell time lapse of wild-type EB1-NG cells carrying out 3 rounds of multiple fission, 3-min. intervals**

Live cell time lapse with 3-min. intervals acquired with microscopy method 1 (see Methods section). Image on left is brightfield, image on right is EB1-NG signal in yellow and chloroplast autofluorescence signal in blue.

**Supp. Video 5. Live cell time lapse of *cycb1-5* EB1-NG cells**

Live cell time lapse acquired with microscopy method 1 (see Methods section), 3 min intervals. Image on left is brightfield, image on right is EB1-NG signal in yellow and chloroplast autofluorescence signal in blue.

**Supp. Video 6. Live cell time lapse of *cdc27-6* EB1-NG cells**

Live cell time lapse acquired with microscopy method 1 (see Methods section), 3 min intervals. Image on left is brightfield, image on right is EB1-NG signal in yellow and chloroplast autofluorescence signal in blue.

**Supp. Video 7. Live cell time lapse of *cdc20-1* EB1-NG cells**

Live cell time lapse acquired with microscopy method 1 (see Methods section), 3 min intervals. Image on left is brightfield, image on right is EB1-NG signal in yellow and chloroplast autofluorescence signal in blue.

**Supp. Video 8. Time lapse microscopy of CYCB1-GFP EB1-mScarlet cells**

Live cell time lapse of CYCB1-GFP EB1-mScarlet cells with 1-min. intervals acquired with microscopy method 3, with 15 z-stacks (see Methods section). Left to right: DIC (time in min:sec indicated); EB1-mSC; CYCB1-GFP; chloroplast autofluorescence; overap of CYCB1-GFP (green) and EB1-mSC (magenta). GFP images were deconvolved to remove chloroplast contribution (Methods). The GFP z images were filtered (0.5 * image (n-1) + image (n) + 0.5* (n-2) and then a maximum projection calculated. This procedure was developed for maximum detail while minimizing graininess. EB1-mSC images are a maximum projection. Control experiments using EB1-mSC cells lacking GFP show no bleedthrough from mSC to the GFP channel (data not shown).

